# *OsLUGL* involved in floral development through regulating auxin level and *OsARFs* expression in rice (*Oryza sative* L.)

**DOI:** 10.1101/552612

**Authors:** C.Y. Yang, D.L. Li, X.J. Zhu, Z.Y. Wei, Z.M. Feng, L. Zhang, J. He, X. Liu, C.L. Mou, L. Jiang, J.M. Wan

**Affiliations:** State Key Laboratory for Crop Genetics and Germplasm Enhancement, Jiangsu Plant Gene Engineering Research Center, Nanjing Agricultural University, Nanjing 210095, China; National Key Facility for Crop Resources and Genetic Improvement, Institute of Crop Science, Chinese Academy of Agricultural Sciences, Beijing 100081, China

**Author notes:** Corresponding authors: Ling Jiang Jianmin Wan.

**Keywords:** rice, OsLUGL, *OsGH*, OsMADSs, floral development, Auxin, *OsARFs*

## Abstract

Specification of floral organ identity is critical for floral morphology and inflorescence architecture. Floral organ identity in plants is controlled by floral homeotic A/B/C/D/E-class genes. Although multiple genes regulate floral organogenesis our understanding of the regulatory network remains fragmentary. Here, we characterized rice floral organ gene *KAIKOUXIAO* (*KKX*), mutation of which produces an uncharacteristic open hull, abnormal seed, and semi-sterility. KKX encodes a putative LEUNIG-like (LUGL) transcriptional co-repressor. *OsLUGL* is preferentially expressed in young panicles and its protein can interact with OsSEU, which functions were reported as an adaptor for LEUNIG. OsLUGL-OsSEU functions together as a transcriptional regulatory complex to control organ identity specification through regulation of MADS-box genes. During this process, SEP3 (such as OsMADS8) and AP1 (such as OsMADS18) serve as the DNA-binding partner of OsLUGL-OsSEU complex. Further studies indicated that OsMADS8 and OsMADS18 could bind the promoter of *OsGH3-8* and regulate its expression. The altered expression of *OsGH3-8* caused the increased auxin level and the decreased expression of *OsARFs*. Overall, our results demonstrate a possible pathway whereby OsLUGL-OsSEU-OsAP1-OsSEP3 complex as a transcriptional regulator by targeting the promoter of *OsGH3-8* directly, affecting auxin level, *OsARFs* expression level and thereby influencing floral development. These findings provide a valuable insight into the molecular functions of OsLUGL in rice floral development.

**Highlight:** OsLUGL through forming OsLUGL-OsSEU-OsAP1-OsSEP3 complex regulate *OsGH3-8* expression, and regulate auxin level and *OsARFs* expression indirectly. This work is a new insight to floral development molecular mechanism.

## Introduction

Rice (*Oryza sativa* L.) as one of the most important cereal crops feeds more than half the world population. Given the rapidly increasing population and decreasing cultivated land area, continued improvements in rice production and quality are massive challenges for rice breeders (Ikeda *et al.*, 2013). Normal development of floral organs in rice is essential for reproduction and seed quality (Zhang *et al.*, 2015). Initiation and differentiation of floral organs are of fundamentally important in the plant life cycle (Zhang *et al.*, 2007; Fornara *et al.*, 2010). A typical dicot flower consists of four whorls, intensive molecular and genetic analyses in *Arabidopsis thaliana* and *Antirrhinum majus*, led to the ABCDE model (Coen and Meyerowitz, 1991; Alvarez-Buylla *et al.*, 2010; Litt and Kramer, 2010). The functions of A/B/C/D/E are to specify the identity of each organ and to control floral meristem determinacy (Coen and Meyerowitz, 1991; Pelaz *et al.*, 2000; Theissen and Saedler, 2001; Pinyopich *et al.*, 2003; Ditta *et al.*, 2004). Rice is a monocot grass species and the model plant for functional genomics studies in crop plants; its spikelet morphogenesis is important for the achievement of yield. Accumulating evidence shows that the ABCDE model is at least partially applicable to floral development in rice (Ferrario *et al.*, 2004; Bommert *et al.*, 2005; Thompson and Hake, 2009; Ciaffi *et al.*, 2011; Tanaka *et al.*, 2013; Zhang and Yuan, 2014; Wang *et al.*, 2015; Dreni and Zhang, 2016;).

Many A/B/C/D/E class genes have been identified, and most of them are MADS-box family genes. Five types of MADS-box genes identified in from *Arabidopsis* are partially conserved in rice, and these genes are reported to be involved in specification of floral development. *OsMADS15* is reported as a regulator of palea size (Wang *et al.*, 2010); *OsMADS16* regulate development of lodicules and stamens (Yun *et al.*, 2013); *OsMADS3* plays a predominant role in stamen specification, whereas *OsMADS58* is involved in establishing floral meristem determinacy and carpel development (Dreni *et al.*, 2011; Hu *et al.*, 2011); *OsMADS13* has a key role in specification of ovule identity and floral meristem determination (Dreni and Kater, 2014; Hu *et al.*, 2015).

Rice has diversified at least five *SEP*-like genes that specify the identities of four-whorl floral organs, such as *OsMADS1*, *OsMADS5*, *OsMADS7*, *OsMADS8*, and *OsMADS34* (Malcomber and Kellogg, 2004, 2005; Zahn *et al.*, 2005; Arora *et al.*, 2007; Wu *et al.*, 2018). Loss-of-function *OsMADS1* shows the outer floret organs, lemma and palea, were narrow, poorly developed, and failed to enclose the inner organs (Khanday *et al.*, 2013). *osmads7* and *osmads8* exhibit late flowering, homeotic transformations of lodicules, stamens, and carpels into palea/lemma-like organs, and a loss of floral determinacy(Cui *et al.*, 2010). *OsMADS6* is *AGL6*-like gene have high sequence similarities with SEP-like genes (Dreni and Zhang, 2016; Callens *et al.*, 2018), in *osmads6*, the palea develops five to six cascular bundles, which resembles the identity of a wild-type lemma (Li *et al.*, 2010; Dreni and Zhang, 2016).

In grass reproductive meristems, the phytohormone auxin plays a central role in almost all developmental and physiological processes, it regulates axillary meristem initiation and outgrowth by controlling cell polarity establishment and cell elongation (Zhao, 2010). Auxin responses are mediated by a class of transcription factors known as *auxin response factors (ARFs)* (Overvoorde *et al.*, 2005; Boer *et al.*, 2014). The functions of *ARFs* are well studied. Many loss-of-function mutations affecting floral morphology have been reported in *Arabidopsis thaliana*. For example, *arf1* and *arf2* affect leaf senescence and floral organ abscission (Ellis *et al.*, 2005); *arf3* displays defects in the gynoecium and floral meristem patterning (Nishimura *et al.*, 2005; Zheng *et al.*, 2018). In rice, the transgenic lines expressing an antisense *OsARF1* showed extremely slow growth, poor vigor, curled leaves, and sterility (Attia *et al.*, 2009). Mutation of *OsARF19* caused abnormal floral organs, and changes to plant height, leaf shape, and seed size (Zhang *et al.*, 2015).

*Arabidopsis* LUG was Gro/Tup1-like co-repressor identified in plants; its role is to regulate transcription of the floral homeotic gene *AGAMOUS* (*AG*) (Conner and Liu, 2000). In loss-of-function mutants of LUG, *AG* is ectopically expressed in the outer two whorls of the flower, converting sepals into carpelloid floral organs and reducing the numbers of petals and stamens (Liu and Meyerowitz, 1995). Like other Gro family members, the N-terminal LUFS domain of LUG is required for repression of transcription and for direct interaction with SEUSS (SEU), SEU acts as an adaptor protein, bridging interaction between LUG and specific DNA-binding transcription factors AP1 and SEP3. The LUG-SEU-AP1-SEP3 complex is directly regulating *AG* expression in all four whorls in *Arabidopsis* (Sridhar *et al.*, 2004; Gregis *et al.*, 2006; Sridhar *et al.*, 2006; Sitaraman *et al.*, 2008; Gregis *et al.*, 2009). There is no report on LUG or its regulatory mechanism in rice.

Here, we characterize rice floral organ gene *KAIKOUXIAO (KKX)*, mutation of which causes an uncharacteristic open glume and abnormal seed. KKX encodes a putative LEUNIG-like (LUGL) transcriptional co-repressor. Further analyses revealed that OsLUGL interacts with OsSEU to become a transcriptional regulated complex. OsSEU also interacts with OsMADS8 and OsMADS18. Also, we confirmed that *OsGH3-8* is the downstream target of OsMADS8 and OsMADS18. In *kkx*, the down-regulated expression of *OsGH3-8* cause high auxin level and altered *OsARFs* expression. Thus, our results suggest that OsLUGL regulates floral organ development by forming OsLUGL-OsSEU-OsAP1-OsSEP3 complex, which could regulate auxin level and signaling directly.

## Materials and methods

### Plant materials

The *kkx* mutant was from a M_2_ population of ^60^Co-irradiated variety Nanjing35 (*Oryza sativa,* L.). The *kkx* was crossed with crossed with the typical *indica* cultivar 93-11 to construct the mapping population. The F_1_ seeds of *kkx*×93-11 were sown and transplanted as individual plants to generate the F_2_ plants for gene mapping. Nanjing35 was used as the wild-type plants for phenotypic analysis. All plants were cultivated in an experimental field under natural long-day conditions in Nanjing, China.

### Fertility evaluation of pollen and embryo sac

Ten individuals from wild-type and *kkx* were examined to determine the pollen fertility. Ten florets from three panicles of each plant were collected 2-3 hours before flowering. All anthers per floret was mixed, placed on the slide, mashed, and stained with 1% iodine potassium iodide (I_2_-KI) solution, and observed with an ECLIPS E80i (Nikon, Tokyo, Japan) light microscope.

To observe the embryo sac development of the wild-type and *kkx*, spikelets were collected immediately after pollens disperse, fixed in FAA solution. Before staining, remove lemma, palea and anthers. The embryo sac was then processed through an ethanol series (70%, 50% and 30%) and finally transferred into distilled water (30 min each). The whole ovary was incubated in 2% aluminum potassium sulfate (KAl (SO_4_)_2_) water solution for 30 min, then held in eosin Y water solution for 10-12 h, and then in 2% aluminum potassium sulfate for 20 min, after washing 2-3 times with distilled water. The ovaries were processed through an ethanol series (30%, 50%, 70%, 80%, 90%, 100% and 100%, 30 min each), then transitted to a mixture of absolute ethyl alcohol and methyl salicylate (1:1) for 1-2 h, the held in methyl salicylate over 12 h. Fertility of embryo sacs was examined by confocal laser scanning microscopy (Leica SP8).

### Microscopy observations

For paraffin sectioning, *kkx* and wild-type flowers from young panicles were fixed in formalin–acetic acid–alcohol (FAA) solution, and samples were treated by a graded series of dehydration and infiltration steps. Fixed tissues were embedded in paraplast. Samples were sectioned to 15 μm, stained with 0.1% toluidine blue and observed with an ECLIPS E80i (Nikon, Tokyo, Japan) light microscope.

For scanning electron microscopy (SEM), young panicles were fixed in 2.5% (v/v) glutaraldehyde, and fixed overnight at 4◻ after vacuuming, and dehydrated through a graded concentration of ethanol. The samples infiltrated and embedded in butyl methyl methacrylate, treated with critical point drying, and then sputter coated with platinum. All tissues were observed with HITACHI S-3000N scanning electron microscope.

### Gene mapping and RNAi suppression of the KKX gene

83 plants with *kkx* phenotypic were selected from the F_2_ mapping population of *kkx* and 93-11 for preliminary mapping using 122 polymorphic SSR (simple sequence repeat) markers between *kkx* and 93-11. Then, 513 F_2_ recessive plants were used for fine mapping. Fine-mapping sequence-tagged site primers were designed according to the different DNA sequences of 93-11 and Nipponbare (*O. sativa, japonica.*) obtained from the National Center for Biotechnology Information (NCBI). Primers used for mapping are listed in Supplementary Table S1.

To obtain *KKX* RNAi plants, the construct LH-1390-RNAi was used as an RNAi vector. Both sense and antisense versions of a specific 200 bp fragment from the cDNA of *KKX* were amplified with primer pairs KKX-RNAi-L and KKX-RNAi-R (Supplementary Table S2), and cloned into LH-1390-RNAi to create the *dsRNAiOsKKX* construct, which was then transformed into the rice variety Nanjing35 by the Agrobacterium-mediated method (Hiei *et al.*, 1994).

### Real-time PCR analysis

Total RNA from seedling, root, shoot, leaf, panicle, young spikelet, and different stage of panicles were isolated using the RNA prep Pure Plant Kit (TIANGEN, Beijing, China). First-strand cDNA was reverse transcribed from 1μg of total RNA using the PrimeScript 1st Strand cDNA Synthesis Kit (TaKaRa). Rice ubiquitin (UBQ) was used as endogenous control. Real-time PCR was performed using a SYBR Premix Ex TaqTM kit (TaKaRa) on an ABI prism 7500 real-time PCR System and three biological repeats. Primers used for real-time PCR analysis are listed in Supplementary Table S3.

### Subcellular localization of LUGL

To explore the subcellular localization of OsLUGL, the C-terminus of OsLUGL cDNA was fused with green fluorescent protein (GFP) and inserted in the pAN580 vector between the d35S promoter and the nopaline synthase (NOS) terminator. In addition, we used the mCherry-tagged rice prolamin box binding facto (RPBF-mCherry) vector as a nuclear marker (Kawakatsu *et al.*, 2009). And the 35s-OsLUGL-GFP plasmid and the nuclear marker plasmid were co-transformed into rice protoplasts. Rice protoplasts were prepared, transfected, and cultured as previously described (Wang *et al.*, 2016).Fluorescence images were observed using a Zeiss LSM510. Primers used to make subcellular localization constructs are listed in Supplementary Table S2.

### RNA in situ hybridization

RNA in situ hybridization was performed as described previously (Chen *et al.*, 2015)with minor modifications. The young panicles of all stages from Nipponbare (*O. sativa, japonica.*) was fixed in FAA solution, subjected to a dehydration series and infiltration, and embedded in paraplast and sectioned at 8μm using a Leica RM2235 microtome. A 360 bp gene-specific region of OsLUGL amplified with primers OsLUGL-PF and OsLUGL-PR (see Supplementary Table S2) was cloned into the pGEM-T Easy vector (Promega). The linearized templates were amplified from the pGEM-T plasmid containing the gene-specific region of SLG using primers Yt7 and Ysp6. Digoxigenin-labeled RNA probes were prepared using a DIG Northern starter kit (Cat. No. 2039672, Roche) following the manufacturer’s instructions. Slides were observed under bright field using a Leica DM5000B microscope.

### Yeast two-hybrid assay

The yeast two-hybrid assays were performed using the Matchmaker Yeast Two-Hybrid System (Clontech). Various fragments from OsLUGL^WT^ and OsLUGL^*kkx*^ were cloned into pGBKT7, and OsSEU was cloned into pGADT7. All constructs were transformed into the recipient strain AH109 and selected on SD/–Trp–Leu (DDO) plates at 30°C for 2-3 days. The interactions were assayed on selective medium SD/-Trp-Leu-His-Ade (QDO) plates at 30°C for 3-5 days. For testing the interaction with OsMADSs, full-length cDNA of OsMADS8/14/15/18 were cloned into pGBKT7, and full-length cDNA of OsMADS6/14/15/18 were cloned into pGADT7. All constructs and/or OsSEU-AD were transformed into the recipient strain AH109 and selected as mentioned above. Primers used to make these construct are listed in Supplementary Table S2.

### Bimolecular fluorescence complementation (BiFC) assay

Full-length cDNA of OsLUGL^WT^ and OsLUGL^*kkx*^ were cloned into the p2YN (eYFP) vector to construct the OsLUGL^WT^ -eYFPN fusion protein and OsLUGL^*kkx*^ -eYFPN fusion protein, respectively. OsSEU was cloned into the p2YC (eYFP) vector to produce OsSEU-eYFPC fusion proteins. Primers used to make these constructs are listed in Supplementary Table S2. These plasmids were co-expressed in tobacco leaf epidermis cells by *Agrobacterium*-mediated infiltration (Hiei *et al.*, 1994). The LJ mCherry ER-rk CD3-959 was used as ER (endoplasmic reticulum) marker (Nelson *et al.*, 2007). Yellow fluorescent protein was observed using a Zeiss LSM510 after 48 h infiltration.

### Transactivation analysis

The full-length OsSEU was fused to the GAL4 DNA BD-coding (Yeast Two-hybrid System, Clontech) sequence and constructed into pAN580, which was cutoff GFP protein, to generate effector plasmid pAN580-GAL4-OsSEU. The full-length OsLUGL^WT^ was fused into the pAN580 (no GFP) to generate effector plasmids pAN580-OsLUGL^WT^. The reporter was a plasmid harbouring firefly LUC (luciferase) gene which was controlled by a modified 35S promoter with 5× the UAS (upstream activating sequence) in it. pRT107 vector containing the BD sequence and BD-VP16 fusion sequence were used as negative and positive control respectively. A pPTRL plasmid that contained a CaMV (Cauliflower mosaic virus) 35S promoter and Renilla LUC, was used as an internal control (Ohta *et al.*, 2000). To test the transactivation of OsMADSs, full-length cDNA of OsMADS8/18 were fused into pAN580 (no GFP) to generate effector. To generate the pOsGH3-8:LUC reporter construct, ~ 2 kb of the OsGH3-8 promoter was cloned into pGreenII-0800-LUC. The different effectors and different reporters were co-transformed into rice protoplasts in different combinations. Rice protoplasts were prepared as mentioned before. The luciferase activity assay was investigated using the dual luciferase reporter assay system and the relative luciferase activity was detected referring to the protocol (Promega, E1910). The primers used for the constructions are listed in Table S2.

### Yeast one-hybrid assay

For yeast one-hybrid assays, the full length cDNA of OsMADS8/18 were cloned into pB42AD vector, and promoter of OsGH3-8 was fused into pLacZi vector. Plasmids were co-transformed into yeast strain EGY48 according to the manufacturer’s manual (Clontech). Transformed yeast was plated onto a synthetic medium DDO (-Ura/-Trp). Positive transformants was screened by adding 80 mg·L^−1^ X-α-gal into SD/-Ura/-Trp medium. The primers used for the constructions are listed in Table S2.

## Results

### Phenotypic characterization of kkx

To investigate the regulation of rice floral development, we identified floral mutant *kkx*, which has displays open hulls, semi-sterile, smaller anthers and pistils, and poor quality seeds (Figure 1a-e). I_2_-KI staining showed normal pollen fertility (Figure 1f, g). Also, observation of the embryo sacs showed that normal embryo sacs in wild-type and *kkx*, of which two polar nuclei located in the central cavity with horizontal arrangement instead of vertical one (Figure S1). In terms of agronomic traits 1,000-grain weight was significantly reduced, and seed set was 57.50±6.40% compared to wild-type at 82.78±6.63%. Plant height and panicle length were similar to wild-type (Figure S1).

**Fig. 1.**
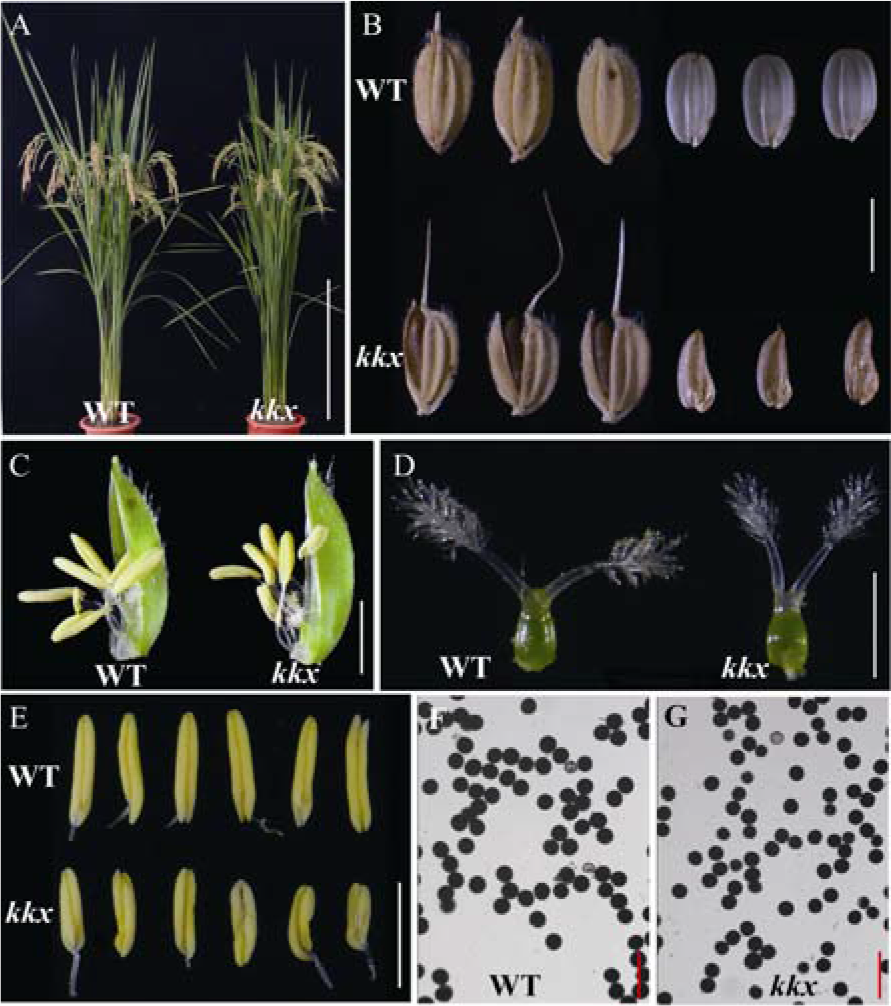
Phenotypic characteristics of wild-type and *kkx*. (A) Phenotype between wild-type and *kkx* at the post-heading stage. Scale bars = 50 cm. (B) Phenotypes of mature seeds with and without seed coat of wild-type and *kkx*. Scale = 5 mm. (C) Spikelets phenotype of wild-type and *kkx* after removal of the palea. Scale bar = 3 mm. (D and E) Comparisons between wild-type and *kkx* in pistils (D) and mature anthers (E). Scale bar = 2 mm. (F and G) Pollen fertility of wild-type and *kkx* observed by staining with 1% I_2_-KI. Scale bar = 600 μm.

### The kkx mutant shows open glumes

The lemma and palea of wild-type florets form an interlocked structure with two hamuli at the marginal regions of the palea (mrp) (Figure 1b and Figure 2a-c). In *kkx,* the abnormally shaped palea cannot interlock with the lemma because of lacking or incomplete hamulus in mrp (Figure 1b and Figure 2d-f). Both wild-type and *kkx* have five vascular bundles in the lemma and three in palea (Figure 2a and d).

**Fig. 2.**
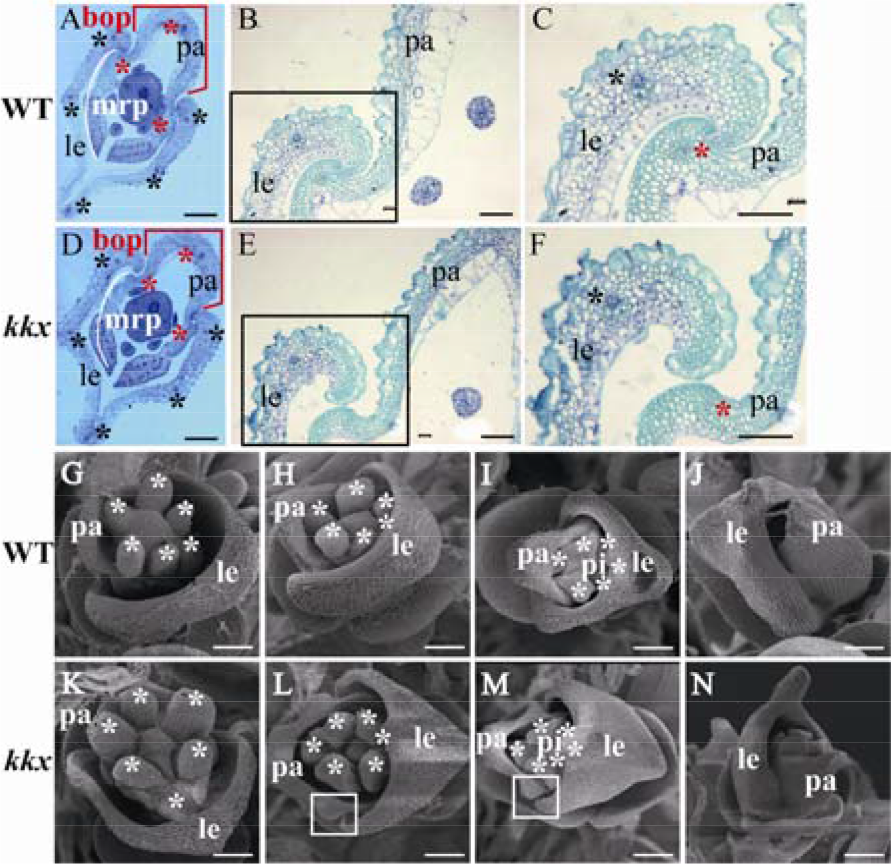
Histological analysis of wild-type and *kkx* spikelets. (A and D) Cross sections of spikelets showing five vascular bundles in the lemma (black asterisks) and three vascular bundles in the palea (red asterisks) in wild-type (A) and *kkx* (D). White lines indicate the mrp of paleas in wild-type (A) and *kkx* (D), red brackets indicate the bop of paleas of wild-type (A) and *kkx* (D). Scale bar = 500 μm. (B-C and E-F) Cross sections of palea edges in wild-type (B and C) and *kkx* (E and F). Black asterisks indicate vascular bundles in the lemma, and red asterisks indicate vascular bundles in the palea. Scale bar =300μm. (G–J) Scanning electron micrographs (SEM) analysis of early spikelets of wild-type at different stages. (G) Sp6; (H) early Sp7; (I) Sp7; (J) Sp8. (K-N) Scanning electron micrographs of early spikelets of *kkx* at different stages. (K) Sp6; (L) early Sp7; (M) Sp7; and (N) Sp8. White square indicate the bulge on palea of *kkx*. White asterisks indicate the stamens in wile-type and *kkx* spikelets, respectively. pi, pistils; le, lemma; pa, palea. Scale bar, 50 μm in (G to N).

We analyzed the early spikelets of wild-type and *kkx* using scanning electron microscopy (SEM). At late stage Sp6 (the stamen primordial formation stage), palea primordia were already formed and the stamen primordia were initiated both in wild-type and *kkx* (Figure 2g and k). At early stage Sp7 and stage Sp7 (pistil primordium formation stage), the pistil primordium formed in the center of the six stamens. The developmental processes of stamen were not significantly different between wild-type and *kxx*, but there was a bulge on palea of *kkx*, which marked with white square (Figure 2h, i, l and m). At stage Sp8 (the ovule and pollen formation stage), wild-type and *kkx* formed a normal spikelet, whereas *kkx* had a larger spacing between the lemma and palea (Figure 2j and n). SEM analysis revealed that both the inner and outer epidermal cells of lemma and palea in *kkx* were similar to those of wild-type (Figure S3).

### OsKKX encodes a putative LEUNIG-Like (LUGL) transcriptional co-repressors

An F_2_ population from a cross of the *kkx* mutant and wild-type segregated 151 individuals with wild-type phenotype and 38 individuals with mutant phenotype, the segregation ratio was consistent with that expected a single locus (χ^2^_3:1_ = 0.418, *P*_*1df*_>0.05). We selected 83 individuals with *kkx* phenotype from the F_2_ progeny of a cross between *kkx* and 93-11 and mapped the locus in a region between InDel marker InDelB and SSR marker RM5496 on the long arm of chromosome 1. Further fine mapping narrowed this region to a 36.8 kb genomic region (Figure 3a and b). According to the Rice Genome Automated Annotation System (RiceGAAS, http://ricegaas.dna.affrc.go.jp) there were three putative genes in this region, coding for one pentatricopeptide protein, and two *LEUNIG-like (LUGL)* transcriptional co-repressors (Figure 3c).

**Fig. 3.**
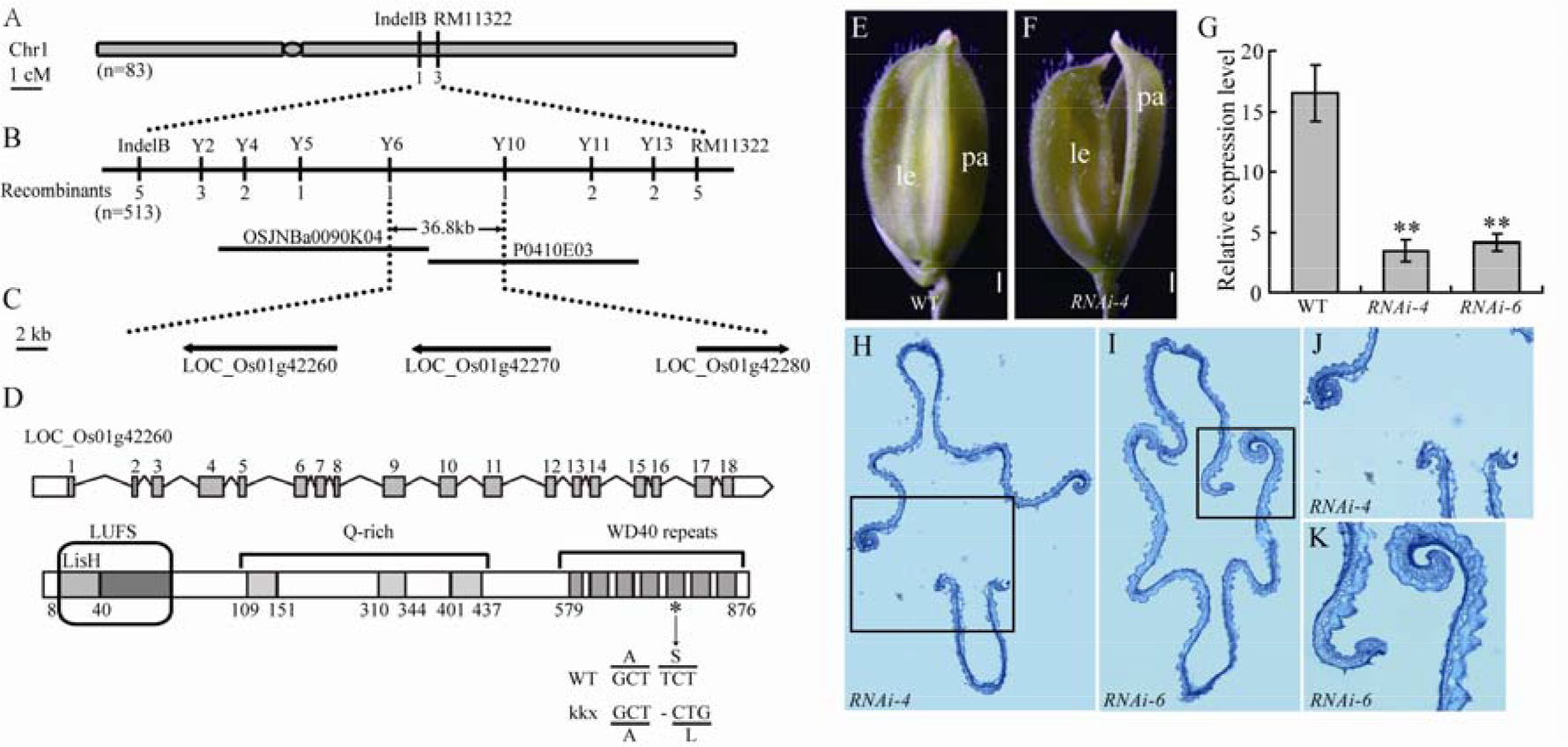
Map-based cloning and identification of *KAIKOUXIAO (KKX)*. (A) *KKX* was preliminarily mapped between markers InterB and RM11322 on chromosome 1. (B) Fine-mapping of the *KKX* locus. *KKX* was positioned on chromosome 1 between BACs OSJNBa0090K04 and P0410E03 within a 36.8 kb region flanked by In-Del markers Y6 and Y10 using 513 mutant individuals. (C) Three predicted open reading frames are located in the 36.8 kb region. (D) Gene structure of candidate gene *LOC_Os01g042260*. Black asterisk and black arrow indicate the position and differing amino acids in wild-type and *kkx*. (E-K) Characterisation of T_0_ transgenic plants. Phenotypes of wild-type (E) and *RNAi* line (F) florets at the heading stage. Scale bar = 0.5 mm. (G) Relative expression levels of *LOC_Os01g042260* in spikelets of wild-type and *RNAi* lines were detected by qRT-PCR, with data normalized to *UBQ* levels. Error bars indicate s.e.m. of the mean of 3 replicates. **, *P* < 0.01, Student’s *t* test. (H-K) Cross sections of *RNAi-4* (H and J) and *RNAi-6* (I and K) flower, respectively.

We chose the two *LEUING* genes, *LOC_Os01g042260* and *LOC_Os01g042270* as potential candidates. Gene expression profiles analyzed on the Rice Expression Profile Database (RiceXPro, http://ricexpro.dna.affrc.go.jp/) indicated that *LOC_Os01g042270* was barely expressed in young inflorescences, and we also could not obtain a PCR product from cDNA extracted from young inflorescences (20 mm) (Figure S4). *LOC_Os01g042260* was choose to further analysis, which encoding a 95.8 kD protein with 876 amino acids, containing three parts, an N-terminal LUFS domain, a central Q-rich domain, and a C-terminal with 7 WD40 repeats (Figure 3d). Sequencing results revealed that the gene in *kkx* had a T deleted in the fifth WD40 repeat, resulting in an S779 to L779 transition and a premature stop codon (Figure 3d, and Figure S5).

To confirm that the deletion in *LOC_Os01g042260* was responsible for the uncharacteristic open floret phenotype of the *kkx* mutant, we generated transgenic plants carrying a *LOC_Os01g042260* RNA interference (*RNAi*) construct. A plasmid containing two 200 bp fragments of wild-type *LOC_Os01g042260* cDNA inserted forward and reverse into an *LH–1390–RNAi* vector was introduced into rice cv. Nanjing 35. Compared to wild-type, positive lines *RNAi-4* and *RNAi-6* showed open florets and had significantly reduced transcripts of *LOC_Os01g042260* (Figure 3e-g). Anatomical observations revealed that lemmas and paleas of *RNAi-4* and *RNAi-6* were unlocked (Figure 4e, h-k). These results confirmed that *LOC_Os01g042260* (*OsLUGL*) was the *KKX* gene.

**Fig. 4.**
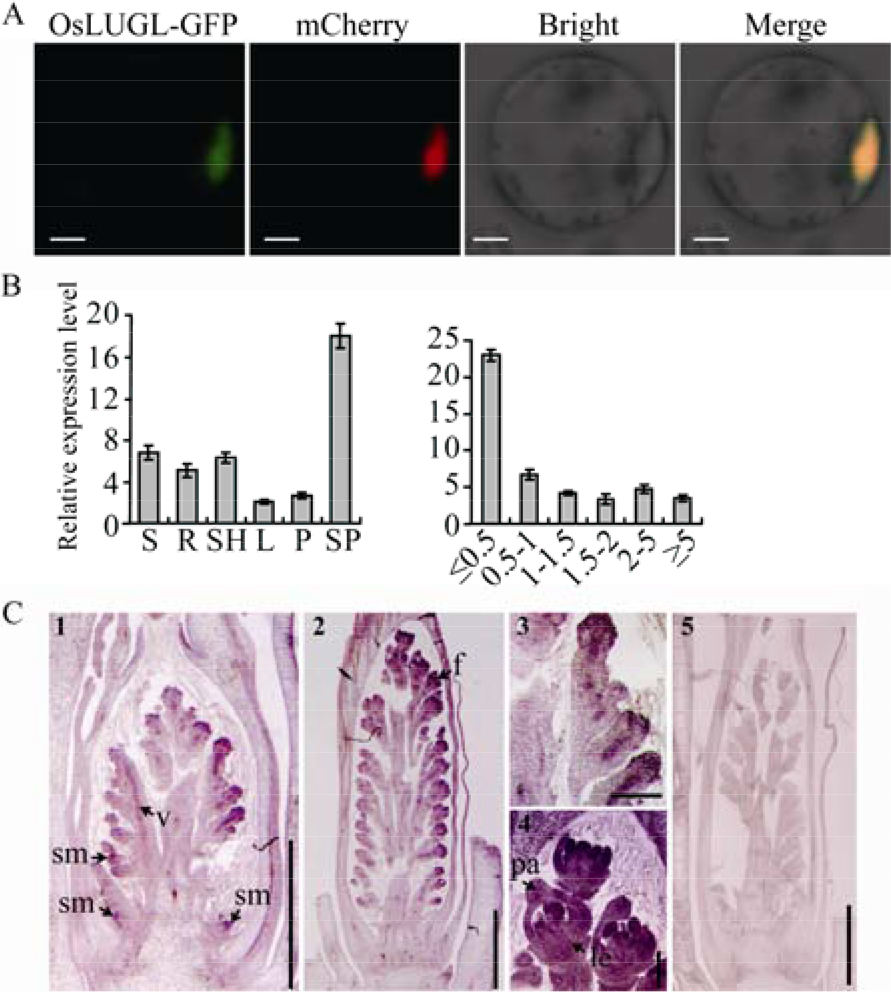
Subcellular localization and expression pattern of OsLUGL. (A) Subcellular localization of OsLUGL protein. Rice protoplast cell expressing OsLUGL-GFP. From left to right: GFP signal of the fusion protein, mCherry with nuclear location gene as a marker, bright field image and merged image. Scale bar, 1 mm. (B) Tissue-specific expression pattern revealed by real-time PCR. RNA was isolated from various tissues of wild-type (lift). S, seedlings; R, roots; SH, shoots; L, leaves; P, panicles; SP, young spikelets. Right image shows the relative mRNA expression of *OsLUGL* in spikelets at different developmental stages. Error bars indicate s.e.m. of the mean of 3 replicates. (C) In situ hybridization of *OsLUGL* mRNA. Panicle development at early (1 and 3) and late (2 and 4) stages of panicle development. Antisense probe was used as the negative control (5).Scale bars, 1 mm in 1, 2 and 5; 100 μm in 3; 200 μm in 4. sm, spikelet meristem; v, vasculature; f, floret; le, lemma; pa, palea.

### OsLUGL was localized in the nucleus and strongly expressed in young panicles

To determine the subcellular localization of OsLUGL, we fused OsLUGL with green fluorescent protein (GFP) at its C-terminus. Transient expression of this fusion protein in rice protoplasts revealed GFP signals in the nucleus (Figure 4a). RT-PCR analysis showed that *OsLUGL* was expressed in all tissues, and strongly in young panicles. Furthermore, *OsLUGL* was most highly expressed during early panicle development, but expression dropped dramatically once the spikelets reached 0.5 cm (Figure 4b). We also performed in-situ hybridization to localize *OsLUGL* expression during early panicle development. Strong expression was detected in spikelet meristem primordia, floral meristem primordia, lemma and palea primordia, and vascular regions (Figure 4c). The results of both RT-PCR analysis and in-situ hybridization implied a role of *OsLUGL* in floral organ development.

### OsLUGL interacts with OsSEU and is required by OsSEU to regulate transcription in planta

Phylogenetic and protein structure analysis found that OsLUGL had high homology, especially the LUFS domain in 15 plants (Figure S6 and S7), and in *Arabidopsis*, the protein At4g32551 was reported that it needs an adaptor protein SEU to form a complex to function as a transcriptional regulator (Sridhar *et al.*, 2004). To investigate the functional forms of OsLUGL, the full-length and different truncated OsLUGL proteins were used for interaction analysis. As shown in Figure 5a, the LUFS domain in the N-terminus of OsLUGL was required for interacting with OsSEU both in wild-type and *kkx*, but Q-rich domain and WD40 repeats were not affect their interaction. And the ability of this interaction was no significantly different between the wild-type and the kkx mutant (Figure 5b). In addition, BiFC analysis also showed that OsLUGL physically interacted with OsSEU in nucleus (Figure 5c).

**Fig. 5.**
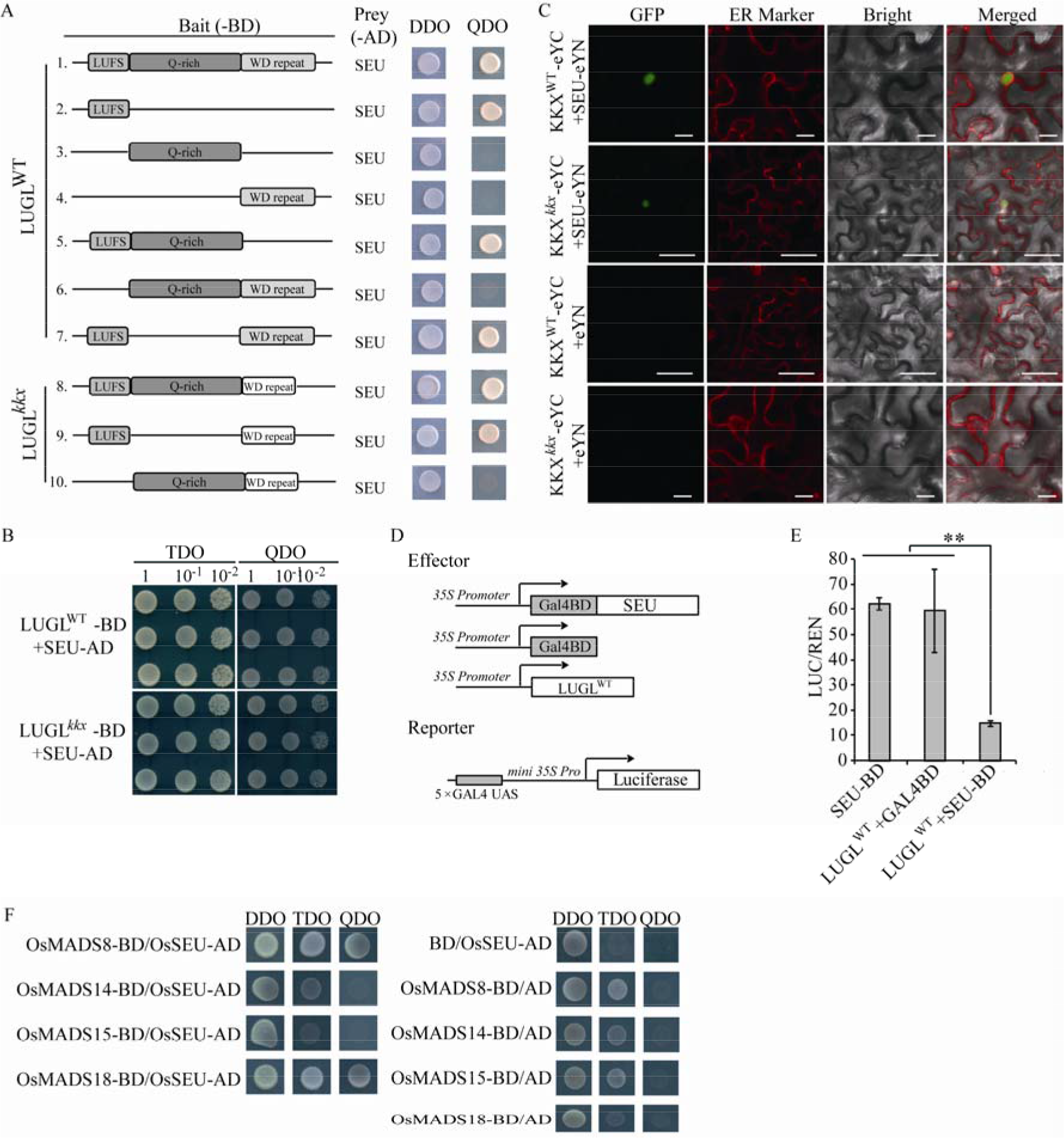
Analysis of LUGL transcriptional regulation mechanism. Yeast two-hybrid assays. Schematic representations indicate the truncated LUGL proteins as baits used for yeast two-hybrid assays. Different colored rectangles represent different domains of LUGL^WT^ and LUGL^*kkx*^ proteins. Yeast diploids were grown on agar plates DDO (-Leu/-Trp) and QDO (-Ade/-His/-Leu/-Trp), respectively. Analysis of interaction strength. Yeast diploids co-expressing Gal4 AD-SEU fusions and GAL4 BD-LUGL^WT^ or GAL4 BD-LUGL^*kkx*^ fusions were grown in selective liquid media to OD600 = 1.0. Diploids were titrated (1, 0.1, 0.01; total 5 μl) and plated on TDO (-His/-Leu/-Trp) and QDO (-Ade/-His/-Leu/-Trp) agar plates for growth. (C) Bimolecular fluorescence complementation (BiFC) assays showing that LUGL interacts with SEU in the nuclei of leaf cells of *Nicotiana benthamiana*. Signals of enhanced yellow fluorescent protein (eYFP) were not detected in corresponding negative controls. ER marker (mCherry ER-rk CD3-959). Scale bar = 20 μm. (D) Diagrams of constructs used for transactivation assays. (E) Transient rice protoplast repression assays using reporter 5×UAS GAL4-LUC mixed with 35S::RenillaLUC (REN). LUC/REN ratio was used to indicate reporter gene expression and to control transfection efficiency. The effectors in (D) were mixed with the reporter to co-transfect rice protoplasts. Data are means ± s.e.m. (n=3). **, *P* < 0.01, Student’s *t* test. (F) Analysis of the interaction between SEU and AP1(OsMADS14/15/18)/SEP1(OsMADS8) genes using yeast two-hybrid assays. Yeast diploids were grown on agar plates DDO (-Leu/-Trp), TDO (-His/-Leu/-Trp) and QDO (-Ade/-His/-Leu/-Trp), respectively.

We next tested whether OsLUGL functioned as a transcription regulator in rice. A transient rice protoplast repression assay was adopted, which 5×UAS GAL4-LUC as reporter. OsSEU was fused to the GAL4 DNA BD-coding sequence and transferred into pAN580 to generate effectors pAN580-SEU (35S::SEU-BD) and using pAN580-GAL4 (35S::GAL4-BD) as a negative control. OsLUGL^WT^ was fused to pAN580 to generate effectors pAN580-LUGL^WT^ (Figure 5d). Rice leaf sheath protoplasts were separately transfected with each plasmid together with the reporter, and LUC expression were quantified. SEU-BD or GAL4-BD alone showed no effect on LUC expression, whereas LUGL^WT^ with SEU-BD significantly reduced LUC expression (Figure 5e). These results confirmed that OsLUGL together with OsSEU functions as a transcriptional regulator in rice.

### OsSEU interacts with SEP3 and AP1 in rice

In *Arabidopsis*, neither LUG nor SEU possesses a recognizable DNA binding motif, the LUG-SEU complex need DNA-binding factor APETALA1 (AP1) and SEPALLATA3 (SEP3) to mediate transcriptional during flower development. During this process, AP1 and SEP3 need to interact with SEU (Sridhar *et al.*, 2006; Sitaraman *et al.*, 2008). Then, we tested whether AP1 and SEP3 serve as the DNA-binding partners of OsLUGL-OsSEU in rice. To determine AP1 or SEP3 that interacts with OsSEU, we performed a yeast two-hybrid assay. OsMADS14, OsMADS15 and OsMADS18 are orthologs of *Arabidopsis* AP1, and OsMADS8 is an ortholog of SEP3 in *Arabidopsis*. OsSEU interacted strongly with OsMADS8 and OsMADS18, but failed to interact with OsMADS14 or OsMADS15 (Figure 5f). These results indicated that OsSEU interact with SEP3 and AP1 in rice.

### Altered expression of floral organ identity genes in kkx

To further understand the function of OsLUGL, we tested the expression of A/B/C/D/E class genes. In real-time PCR analysis, A-class (*OsMADS14/18*), B-class (*OsMADS2* and *OsMADS16*), C-class (*OsMADS3/58*) were down-regulated in *kkx* inflorescences (15 mm); A-class (*OsMADS15*) was up-regulated; E-class (*OsMADS7/8/34/57*) genes were down-regulated; E-class (*OsMADS22/29* and *OsCFO1*) were up-regulated. C-class (*OsDL*) gene, D-class (*OsMADS13*) gene and E-class (*OsMADS1/6*) genes did not vary (Figure 6). *REP1* is a *CYCLOIDEA*-like gene, *osrep1* shows defects in the palea, but not in lemma (Yuan *et al.*, 2009). In *kkx*, the expression of *OsREP1* was down-regulated (Figure 6a). These results showed that the OsLUGL could regulate the expression of floral organ identity genes in rice.

**Fig. 6.**
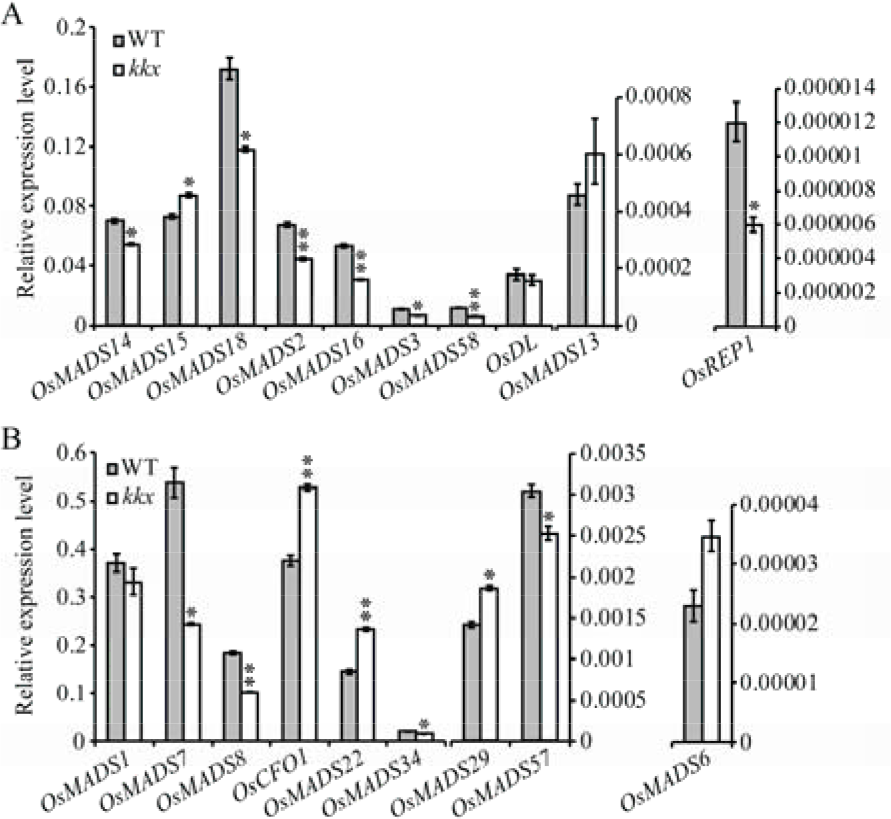
The expression level of floral organ development genes in young inflorescences (15 mm) of wild-type and *kkx*, respectively. (A) Expression of A-class genes (*OsMADS14, OsMADS15* and *OsMADS18*), B-class genes (*OsMADS2* and *OsMADS16*), C-class genes (*OsMADS3, OsMADS58* and *DL*), D-class genes (*OsMADS13*) and *REP1*. (B) Expression of E-class genes (*OsMADS1*, *OsMADS6*, *OsMADS7*, *OsMADS8*, *OsCFO1*, *OsMADS22*, *OsMADS34*, *OsMADS29* and *OsMADS57*). Data are given as means ± s.e.m. (n=3). * and **, *P*<0.05 and 0.01, respectively, Student’s *t* test.

### OsLUGL-OsSEU-OsAP1-OsSEP3 complex has regulatory effects on OsGH3-8 expression, auxin level and auxin signaling pathways

The phytohormone auxin plays a central role in almost all developmental processes, it regulates axillary meristem initiation (Zhao, 2010; Habets and Offringa, 2014). *OsGH3-8* (*OsMGH3*) is an auxin negative component in regulating auxin level and activity, *dsRNAiOsMGH3* caused phenotypes consistent with auxin overproduction or activated signaling, such as ectopic rooting from aerial nodes, carpel development, glume form, pollen viability and reduced fertility (Yadav *et al.*, 2011), which similar to the *kkx* mutant. Also, OsMADS6 is positive regulator for *OsGH3-8* (Prasad *et al.*, 2005; Zhang *et al.*, 2010; Yadav *et al.*, 2011) (Figure 7a and b). In *kkx*, the expression of *OsGH3-8* was down-regulated, but no change in *OsMADS6*, so, a question arises here. What causes the down-regulated of *OsGH3-8*? In *Arabidopsis*, neither OsLUGL nor OsSEU regulate genes expression directly, because the absent of DNA-binding domain. The LUG-SEU complex needs AP1 and SEP3 act as bridge to binding DNA downstream. In this study, we confirmed that OsSEU can interact with OsMADS8 (SEP3) and OsMADS18 (AP1) (Figure 5f), and OsMADS8 and OsMADS18 have high homogeneous of OsMADS6, so we suspect that *OsGH3-8* is direct target of OsMADS8 and/or OsMADS18, OsLUGL-OsSEU-OsAP1-OsSEP3 complex could regulating OsGH3-8 directly. To test this hypothesis, we performed yeast one hybrid assays using in vitro-expressed OsMADA8 and OsMADS18. As shown in Figure 7a, both of OsMADS8 and OsMADS18 bound to the *OsGH3-8* promoter. To examine the regulation of OsMADS8/18 on the expression of *OsGH3-8*, as shown in Figure 7b, we performed transient expression assays using ~ 2 kb of the *OsGH3-8* promoter fused with LUC as a reporter, and OsMADS8/18 were expressed under control of 35S promoter as effectors. Various effectors and reporter were transfect together into rice protoplasts in different combinations (Figure 7c). Higher LUC activity was detected when OsMADS8/18 protein was transfected with the reporter construct compared with the internal control. Meanwhile, we found that the LUC activity is higher in which when two effectors (OsMADS8 and OsMADS18) co-transfected with pOsMGH3-8 than that when single effector (OsMADS8 or OsMADS18) co-transfected with pOsMGH3-8. These findings strongly support the hypothesis that OsMADS8 and OsMADS18 regulate *OsGH3-8* by binding the promoter of *OsGH3-8*, and OsMADS8 and OsMADS18 have a superposition effect on regulation.

**Fig. 7.**
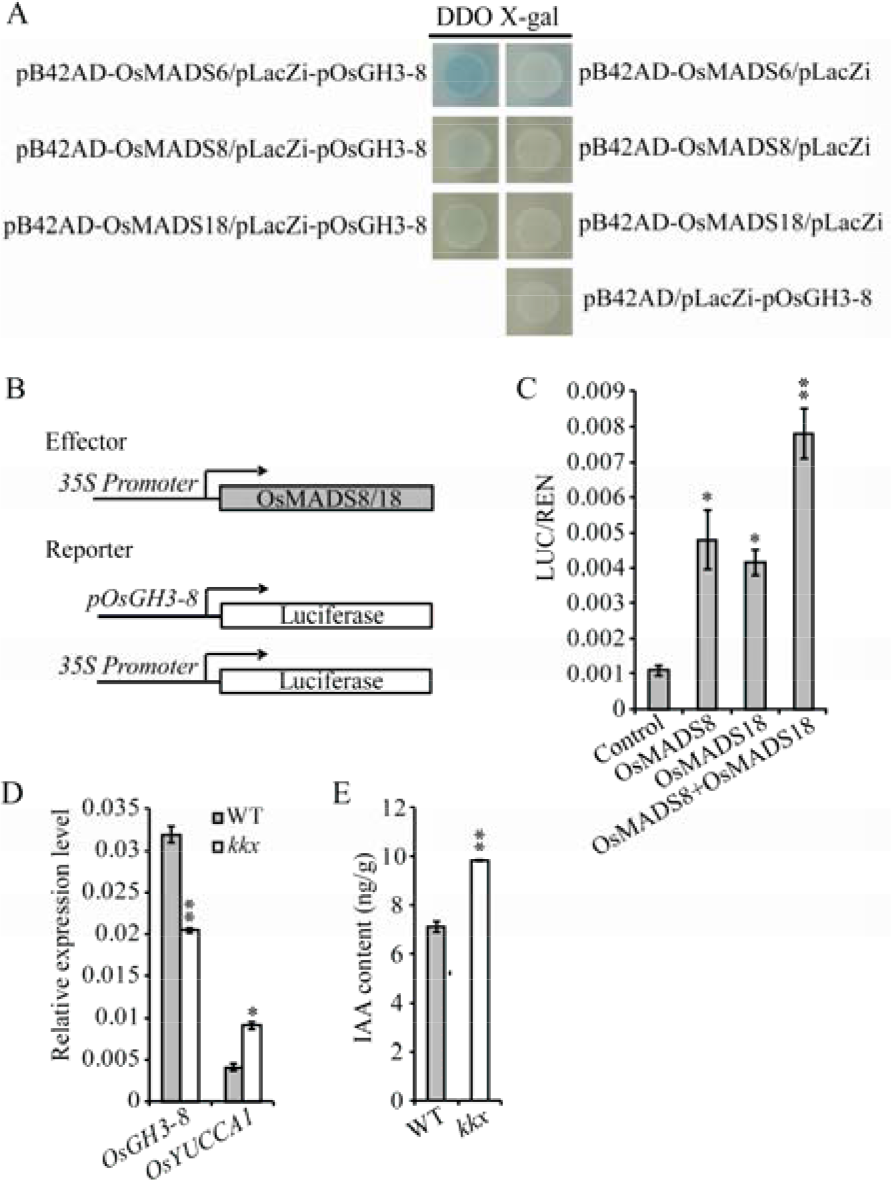
OsMADSs promote the expression of *OsGH3-8* and change the level of auxin level. (A) Yeast one-hybrid (Y1H) analysis of OsMADS6, OsMADS8, OsMADS18 and *OsGH3-8* promoter., *OsGH3-8* as the bait vector which promoter fragment-fused lacZ reporter, and the prey vectors containing OsMADS6/8/18-fused GAL1 activation domain. Two vectors were co-transformed into EGY48. (B) Diagrams of constructs used for transactivation assays. (C) Transient expression assays of *OsGH3-8* transcriptional activity modulated by OsMADS8, OsMADS18 and OsMADS8+OsMADS18 respectively in rice protoplasts. *pOsGH3-8*:LUC was co-transformed with either the effector or empty vector, as control, into rice protoplasts.. LUC/REN indicating the level of *OsGH3-8* expression activated by the effectors mentioned before. (D) Expression of IAA relative genes. (E) The level of auxin in young inflorescences (15 mm). Data in (C-E) are given as means ± s.e.m. (n=3). * and **, P<0.05 and 0.01, respectively, Student’s *t* test.

The phenotypes of *dsRNAiOsMGH3* are similar to *OsYUCCA* overexpression transgenic rice. *OsYUCCA* is a rice auxin biosynthetic gene, and the overproduction of auxin in its overexpression transgenic lines is expected (Yamamoto *et al.*, 2007). In *kkx*, we found the expression of *OsYUCCA1* is up-regulated, and raised auxin level (Figure 7d and e). We also analyzed the expression of all *OsARFs* in young inflorescences (15 mm). The results show that expressions of almost all *OsARFs* were reduced in *kkx*, except *OsARF8, OsARF10, OsARF15,* and *OsARF20* (Figure S9a-d), which suggested that OsLUGL affected floral organ formation and development by regulating the auxin level and signaling pathway.

## Discussion

Rice is model plant for functional genomics studies in crop plants, its spikelet morphogenesis is important to the achievement of yield. In this study, we identified a mutant *kkx* with unlocked hull phenotype, reduced fertility and bugle in Sp7 palea (Figure 1b and 2m), which indicated that the function of *KKX* in spikelet development. We confirmed that the *KKX* encoded a LUG-like transcriptional regulator. LUG, a member of the Groucho family in *Arabidopsis*, acts as a transcriptional co-repressor to regulate plant development and hormonal signaling (Liu and Meyerowitz, 1995; Conner and Liu, 2000; Sitaraman *et al.*, 2008; Grigorova *et al.*, 2011). LUG to require adaptor protein SEU to form a LUG-SEU complex in order to regulating gene expression (Sridhar *et al.*, 2004). Neither LUG nor SEU possesses a recognizable DNA binding motif, they need AP1 and SEP3 to act as the DNA-binding partners (Gregis *et al.*, 2006; Sridhar *et al.*, 2006; Gregis *et al.*, 2009). Our study demonstrated that OsLUGL also needs to interact with OsSEU to form a complex that acts as a transcriptional regulator (Figure 5a-e), and OsSEU could interacted with SEP3 (OsMADS8) and AP1 (OsMADS18) (Figure 5f) to form OsLUGL-OsSEU-OsAP1-OsSEP3 complex in rice.

Expression analysis of some floral organ-related genes in wild-type and *kkx* showed that *OsLUGL* acted as a positive regulator of almost all A/B/C/D/E genes. In *Arabidopsis*, there is a ABCDE model of floral organ specification, A-class genes specify the identity of sepals in whorl 1; A- and B-class genes function together determine the identity of petals in whorl 2; B- and C-class genes coordinated define stamen identity in whorl 3; and C- and D-class genes act to specify carpels in whorl 4. E-class genes are co-regulator with A-, B-, C- and D-class genes during floral identity in all whorls (Coen and Meyerowitz, 1991; Alvarez-Buylla *et al.*, 2010; Litt and Kramer, 2010). Previous research has shown that A-class genes and C-class genes function antagonize each other, and LUG represses the expression of *AG* genes (C-class) (Ma and dePamphilis, 2000). There is a similar ABCDE model in rice (Ferrario *et al.*, 2004; Bommert *et al.*, 2005; Thompson and Hake, 2009; Ciaffi *et al.*, 2011; Tanaka *et al.*, 2013; Zhang and Yuan, 2014; Wang *et al.*, 2015; Dreni and Zhang, 2016). But, in this study, in *kkx*, A-class genes and C-class genes are all down-regulated. We also test their expression in 5 mm young inflorescences, the expression of A-class and C-class genes still down-regulated (Figure S10). These results indicated that OsLUGL-OsSEU might have different role in regulating A/B/C/D gene expression in rice, or there are another unknown factors involved in this regulated process.

Previous reports considered that normal development of mrp may require *OsMADS1* and *OsMADS6* (Khanday et al., 2013; Tao et al., 2018). Mutation of *OsMADS1* resulted in uncharacteristic flowers and/or loss of flower determinacy, suggesting a role of *OsMADS1* in specifying determinacy of the flower meristem, and effects on development of all floral organs (Hu *et al.*, 2015). An *osmads6* mutant showed defects in palea identity; the palea was half open and likely caused by lack of interlocking between lemma and palea (Ohmori *et al.*, 2009; Li *et al.*, 2010; Tao *et al.*, 2018). Similarly, in *kkx*, the glume was half opened (Figure 1b). Compared with *osmads1* and *osmads6*, the phenotypes of *kkx* were weaker than *osmads1* and similar to *osmads6*. Confusingly, the expression of *OsMADS1* and *OsMADS6* were no significant different between wild-type and *kkx* (Figure 6a-c). So, we suspect there might be another factors or pathway to regulating the floral development in rice. *OsGH3-8* is a downstream target of OsMADS6 and an auxin negative component in regulating auxin level and activity, and *dsRNAiOsMGH3* line shows the similar phenotype as *kkx* (Yadav *et al.*, 2011). In this study, we confirmed that OsMADS8 and OsMADS18 can act as a positive regulator of *OsGH3-8* (Figure 7). Also, the expression of *OsMADS8*, *OsMADS18* and *OsGH3-8* were repressed in *kkx* (Figure 6 and 7d). There are 12 members of *GH* in rice, we test the other *OsGH* in *kkx*, most of them were down-regulated except *OsGH3-12* and *OsGH3-13*, but the expression of *OsGH3-12* and *OsGH3-13* were very low (Figure S11). The repressed transcriptional of *OsGH* indicate that OsMADS8 and/or OsMADS18, or the other OsMADS-box genes could regulate *OsGH* expression directly, which need to be further studied. All these results show that OsLUGL-OsSEU-OsAP1-OsSEP3 could regulate the *OsGH* transcriptional directly. But, in this study, the reasons for the altered expression of OsMADS8 and OsMADS8 are still unknown. Besides, the level of auxin in *kkx* was increased, which indicate that OsLUGL affect the auxin level, and OsLUGL regulate floral development by auxin relative pathway.

Morphogen-like properties in developmental processes, such as meristem specification and lateral organ formation, are often attributed to auxin. These processes are mediated by Aux/IAA induced protein degradation and subsequent *auxin response factor (ARF)* activation (Guilfoyle and Hagen, 2007; Zhang and Yuan, 2014; Weijers and Wagner, 2016). Many *ARFs* in *Arabidopsis*, such as *arf1*, *arf2*, and *arf3* are reported to affect floral organs (Ellis *et al.*, 2005; Nishimura *et al.*, 2005; Zheng *et al.*, 2018). We described rice mutant, *osarf19*, in which florets displayed three types of abnormalities. The first was an additional lemma-like organ on the same side as the palea, the second was an enlarged palea with a curved tip, which generated an unclosed floret, and the third was a variably degenerated palea (Zhang *et al.*, 2015). In the present study, we assayed expression of *OsARFs* and most of them were down-regulated in *kkx* (Figure S9). This result provides the evidence that the auxin-dependent gene expression relies on the inhibiting role of auxin, as inhibitors of the *ARFs*. So we confirm that OsLUGL regulate floral development by regulate the *OsARFs* expression.

In previous study, OsMADS6 interact with OsMADS13, and redundantly regulate carpel/ovule identity and floral determinacy (Li *et al.*, 2011). OsMADS6 was also shown to physically interact with OsMADS4 during early flower development (Seok *et al.*, 2010). Also, in rice flower development, OsMADS1 interact with OsMADS6, and B-, C- and D-class proteins (Hu *et al.*, 2015). So, we suspect that AP1 protein interact with SEP3 protein. We test the interaction of OsMADS8 (SEP3 gene), OsMADS14, OSMADS15 and OsMADS18 (AP1 genes). Yeast two-hybrid assays demonstrated that OsMADS8 physically interacts with OsMADS14, OsMADS15 and OsMADS18, respectively. Also, OsMADS14, OsMADs15 and OsMADS18 could interact with each other in yeast (Figure S12). This result indicated that SEP3 and AP1 could co-regulate the expression of OsGH3-8 by form a complex. And it provides the evidence that the OsMADS8 and OSMADS18 have a superposition effect on regulating *OsGH3-8* (Figure 7c).

Collectively, our findings show that OsLUGL acts like a transcriptional regulator in regulating the expression of A-, B-, C-, D-, and E-class genes; OsLUGL-OsSEU-OsAP1-OsSEP3 complex could participate in regulation of flower development by regulation *OsGH* expression directly, and indirectly regulating the auxin level and *OsARFs* expression. SEP3 and AP1 worked together to act as co-regulator in regulate *OsGH3-8* expression. However, there are still unanswered questions. We have no direct evidence to explain the regulate relationship between OsLUGL-OsSEU and *OsMADS8/18*. Why there is no transcription repress in this work? We still have no evidence to explain this issue.

Normal floral structure is necessary for seed production and quality. An abnormal floret, even a half-opened hull, with normal pollen fertility causes a significant reduction in seed setting because the opened glume cannot prevent rain or dew from entering the floret during the fertilization and early grain-filling stages. Also, an abnormal floret will not provide an optimum space for seed development, thus causing smaller, malformed seeds with poor color and quality. Hence, investigation of the regulatory mechanisms underlying floral development is necessary for rice breeding and production.

## Acknowledgments

We thank the Key Laboratory of Biology, Genetics and Breeding of Japonica Rice in the Mid-lower Yangtze River, Ministry of Agriculture, P.R. China, and Jiangsu Collaborative Innovation Center for Modern Crop Production for support. This research was supported by the National Key Research and Development Program of China (2017YFD0100401), National Key Transform Program (2016ZX08001004) and Jiangsu Science and Technology Development Program (BE2018388).

## Supplementary data

Figure S1. Observation of embryo sac development.

Figure S2. Agronomic traits contrast between wild-type and *kkx*.

Figure S3. Scanning electron micrographs (SEM) analysis epidermis cells of lemma and palea in wild-type and *kkx*.

Figure S4. Gene expression profile of *LOC_Os01g042270*.

Figure S5. Alignment of nucleic acid sequence and protein sequence of *LOC_Os01g042260*.

Figure S6. Phylogenetic analysis of the LUGL family in 15 plants.

Figure S7. The amino acid homology sequence analysis of KKX.

Figure S8. The amino acid homology sequence analysis of OsMADS6, OsMADS8 and OsMADS18.

Figure S9. Expression levels of rice *ARF* family genes in wild-type and *kkx* young inflorescences (15 mm).

Figure S10. Expression levels of floral organ development genes in young inflorescences (5 mm) of wild-type and *kkx*, respectively.

Figure S11. Expression levels of rice *GH* family genes in wild-type and *kkx* young inflorescences (15 mm).

Figure S12. Analysis of the interaction between OsMADSs genes using yeast two-hybrid assays.

Table S1. Primers used in mapping.

Table S2. Primers used for vector construction in this study. Table S3. Primers used for real-time PCR analysis.

